# *Fossil-explorer.com*: An efficient interactive approach to exploring fossil data

**DOI:** 10.1101/2022.06.12.495851

**Authors:** Junqi Wu, Honghe Xu, Zhaohui Pan, Zhibin Niu

## Abstract

Fossils today are increasingly being digitized and documented by multi-modal data obtained from visual data (i.e., photos and tomographic images), as well as text, locations, geological ages, and other chemical and physical measurements. Popular online websites such as PBDB and GBDB offer visual explorations of specimens’ localities, but they have limited multi-modal data visualization abilities and face challenges related to visual obscuration and insufficient interaction/exploration. Here, we present fossil-explorer.com, a continuously developing open-source online tool for assisting paleontologists with interactively exploring fossil collections. The tool is designed to address the issues of visual clutter, limited data types, and insufficient interactions. It is intuitive and endorsed by paleontologists. We have also quantitatively evaluated the tool by measuring the interaction scaling performance. The results show that it provides *sublinear interaction performance* and thus is able to deal efficiently with millions-level data. The current `fossil-explorer.com` demonstrates the Ordovician to Silurian graptolite fossil multimedia dataset, which is significant in global stratigraphy and shale gas exploration. The extended version also facilitates the use of Deepbone (http://deepbone.org), world’s most comprehensive database of vertebrate paleontology database. We developed the code for `fossil-explorer.com` to be open access and will continue to improve it.

## 1 INTRODUCTION

Databases are key for breaking down paleontology disciplinary barriers and conducting interdisciplinary research [7]. Fossils are usually digitally archived in a hybrid data form that is both structured (localities, ages, reference, etc.) and unstructured (text, photos, tomographic images, chemical or physical measurements, etc.). These data were traditionally locally stored by category inside paleontologists’ computers. This arrangement is suitable for paleontologic study but makes sharing with others challenging and incredibly inefficient, especially when conducting evolution analyses, due to the large volume of information. Currently, paleontologists prefer to create paleontology databases with paleo/neo-map-based visualization web services to assist with data sharing and big science questions pursued via data-driven approaches [4, 18].

Mainstream paleontology databases systems usually provide web-based fossil records visualization and filtering functions [18]. They might support limited data types and face challenges of visual occlusion and performance issues. Figure 1 shows the two most significant paleontology databases (PBDB and GBDB). It can be seen that there still exists serious visual occlusion. This is because fossils could be collected in the same or different locations (geospatial co-occurrence) and rock layers (geological time co-occurrence). The vast number of fossils make projections overcrowded on maps. Mainstream paleontology databases usually adopt the visual aggregation method (size of marker point) to reduce the visual clutters. Users have to examine the detail list by clicking the nodes. It is *not intuitive, inefficient*, and thus, might discourage the potential of in-depth data sense-making. In addition, the interaction performance is still not fully satisfied (Figure 3). The number of rendered projections imposes a severe performance burden, making interactions not so smooth. These practical issues require us to consider a more effective and efficient visualization method.

**Figure 1:**
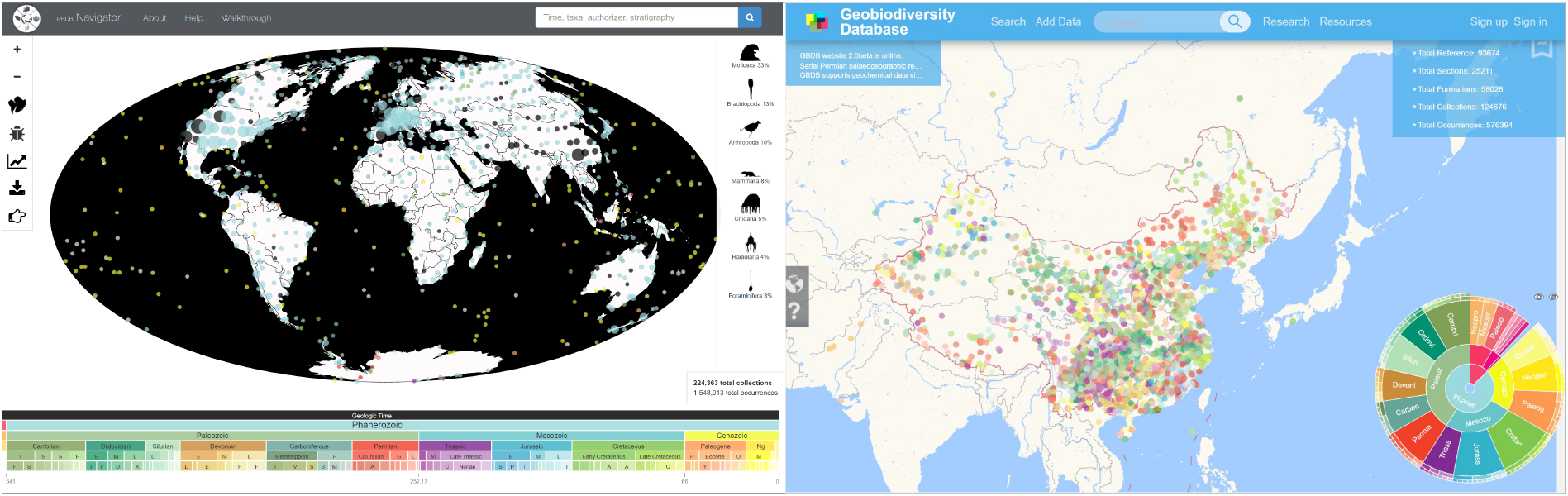
PBDB and GBDB screenshots (March 21st, 2022).

In this research, we describe the prototype for and demonstration of `fossil-explorer.com`. The tool is designed to assist paleontologists with interactively exploring fossil data and address issues related to visual clutter/occlusion, limited data types, and insufficient interactions. Specifically, (i) we designed an intuitive fossil lens glyph to mitigate the visual occlusion issue and (ii) an elaborate technology stack that assures sublinear interaction performance. Scaling experiments showed that our prototype provides smooth interactions at millions-level point projections. Through the visualization of `fossil-explorer.com`, we have released an Ordovician to Silurian graptolite fossil multimedia dataset that will be significant for global stratigraphy and shale gas exploration [20]. The extended version also facilitates the use of Deepbone (http://deepbone.org), the world’s largest vertebrate paleontology database [15]. We personally designed the code for `fossil-explorer.com` to be open access and will continue to improve it.

## 2 PROBLEM STATEMENT

Fossils are collected during field studies of the earth’s surface or through deep-well drilling. The digitalized information usually has the following attributes.

### Taxonomic rank

The fossils are named according to a hierarchy ranging from finer to grosser, as follows: species, genus, family, order, class, phylum, kingdom, and domain. Actual fossil species numbers can be huge. Fossil biodiversity research has estimated that even considering the incompleteness of the record, there are over 5 billion species that have gone extinct over the past 4.5 million years of evolution [1, 8, 14].

### Geological time

The geological timescale is defined through the various layers of rock (i.e., stratigraphy) [5]. Curated fossils are usually linked to a specific geological timescale, which also has a hierarchical structure. For example, the age of the earth can be defined using eons, eras, and periods. The Paleozoic Era includes six periods: Permian, Carboniferous, Devonian, Silurian, Ordovician, and Cambrian.

### Locality

Each fossil has an excavation site localized by GPS and may have a PaleoGPS generated by an algorithm. Note that many fossils can be collected in a sample location but belong to different taxa or different rock layers.

### Multimedia

Paleontologists use photographs and tomographic images or videos to observe and share fine details of fossils.

### Miscellaneous

Other unstructured data are also essential for digital archiving. For example, paleontologists may keep textual descriptions of fossil collections, geochemical indicators, physical measurements, related publications, and other data.

Mainstream paleontology databases provide recognized interactive exploration abilities; however, they still face the following challenges.

### C.1. Serious visual clutters/occlusion

Various taxa fossils can be collected from the same or different locations (geospatial cooccurrence, rock layers or geological timescale co-occurrence). The vast number of fossils makes projections visually overcrowded on maps. Mainstream paleontology databases such as PBDB and GBDB adopt the visual aggregation method (i.e., the size of the marker points) to reduce visual clutter. Users must examine the detail list by clicking on a node. The process is unintuitive and inefficient, and therefore may discourage in-depth interpretation.

### C.2. Limited data types

There is an increasing trend in which more and more forms of fossil data are openly available online for research and dissemination. However, many sources, including PBDB and GBDB, currently provide only fossil record data. Others choose to use a multimedia gallery as the entrance ^1^. This is suitable for exhibition but lacks large-scale geo-spatiotemporal cross-taxa visualization and analytics abilities.

### C.3. Insufficient performance

The interactive visualization of fossil data faces substantial challenges with regards to scaling. The greater body of fossil data is increasing [13] due to the Deep-Time Digital Earth Program [18]. Mainstream paleontology databases provide hundreds of thousands of lines of data that are visualized but allow for limited interaction. In the future, fossil databases may require millions or even billions-level visualization abilities. Many rendered projections impose severe visualization and interaction performance burdens. Unsatisfactory visualization performance makes rich interactions impossible and consequently inhibits interactive knowledge mining.

## 3 THE APPROACH

We populated `fossil-explorer.com` with the Ordovician to Silurian graptolite fossil multimedia dataset [20] (see Figure 2).

**Figure 2:**
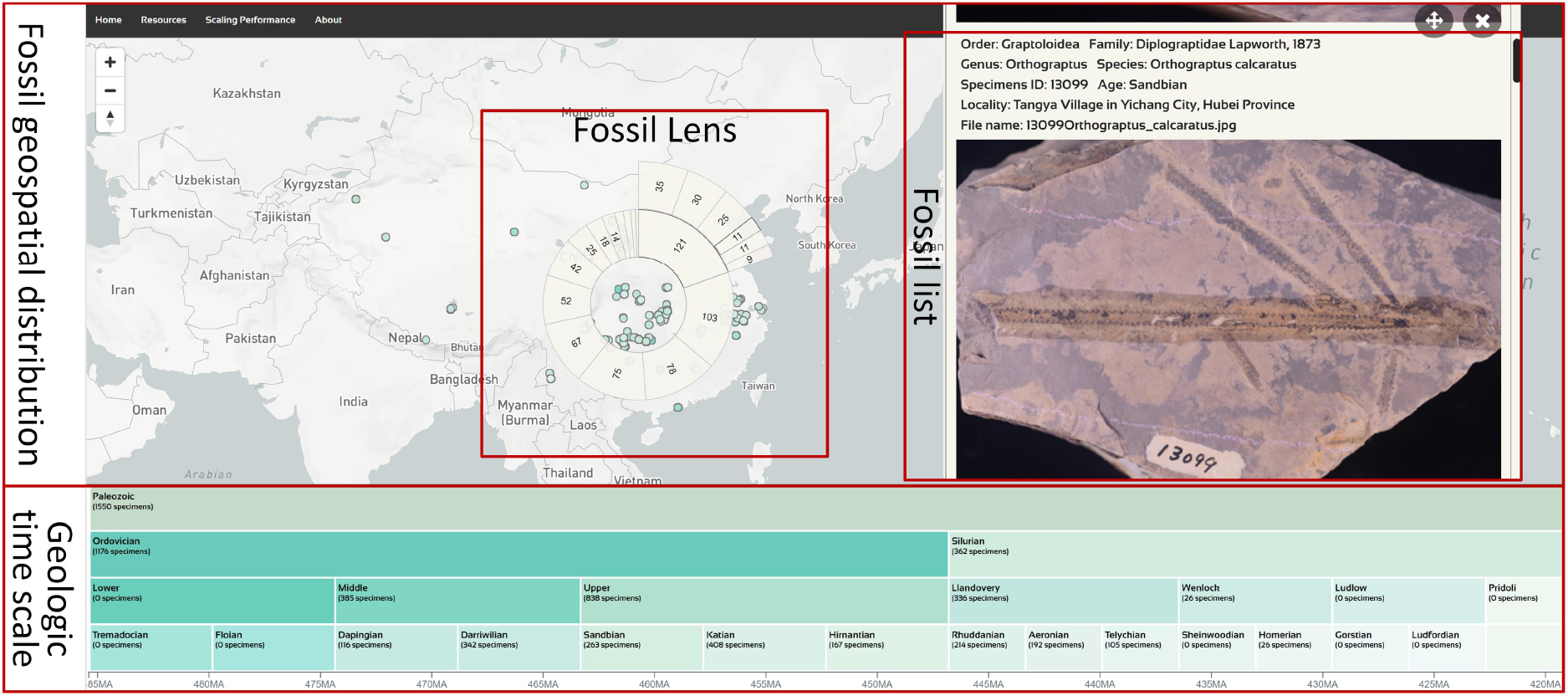
System interface for Fossil-explorer.com.

**Figure 3:**
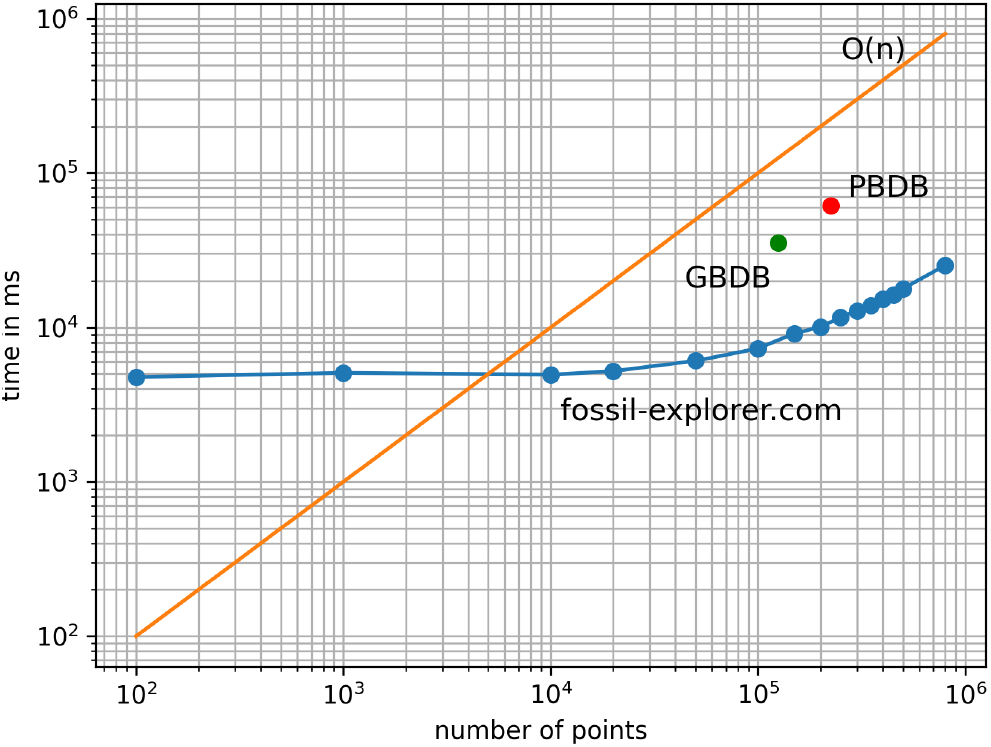
The interaction performance of PBDB, GBDB, and `fossil-explorer.com`. The PBDB has 223,831 fossil collections, and the interactions take 61.6 seconds. The GBDB has 124,675 fossil collections, and the interactions take 35 seconds. Our `fossil-explorer.com` has much better interaction performance; even for 800,000 fossil collections, it only takes 25.3 seconds for all interactions.

There are two main coordinates needed for interactive explorative analysis. **Geological timescale view** 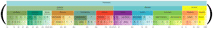 uses the standard geological timescale employed in paleontology [5], which is technologically charted as a tree map chart with filtering ability. We also give the aggraded number of specimens for each timescale. Similar to PBDB and GBDB, we chose geospatial localization as the starting point. The **fossil geospatial distribution view** projects the fossil collection locations; the **fossil lens** is a tailor-made specimen selector that allows users to observe fossils from a particular region. The inner and outer rings represent the family and genus. When the user chooses a specific genus, the **fossil list view** gives details.

To reduce the visual occlusion of overcrowded fossil data (C.1), we designed **fossil lens** 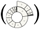 to exploit the hierarchy of taxonomic ranks. Fossil lens follows mouse movement and then aggregates and visualizes the fossil data in that area. In our test case, the circular tree map includes two layers: family and genus. For more general use, it can represent the whole taxonomic rank. The sizes of the tree map cells are in proportion to the number of taxa types. *The design for fossil lens avoids visual occlusion and is able to chart the biodiversity of the selected area*.

Fossil lens interactions allow users to view record details (C.2, C.3). At first, it only visualizes the first layer. After double clicking on the detector, it becomes fixed and no longer follows the mouse movement. Users can click the circular tree map cell to toggle to the second layer. When mousing over the cell of the circular tree map, the tool tip shows the upper taxon name and genus sub taxon, along with their numerical magnitudes. Clicking on the circular tree map cell of the genus layer triggers the **fossil list view**. Double-clicking again on the fossil lens resets it and activates the following function.

### Fossil list view

This function displays detailed fossil information with multimedia data. In our current demonstration, each record consists of the taxonomic rank and fossil image. Each image on the list is bound to a function to trigger the fully functional image viewer. Scrolling allows the user to view other selected records when the mouse is focused on that list. The close button closes the fossil list, while the drag button moves the list window to anywhere that is convenient for observation. Other data types such as chemical or physical measurements and additional multimedia data such as tomography can be appended to the fossil list view.

#### The interaction and scaling performance is also a bottleneck and challenge

As explained in C.3, current mainstream fossil collection systems usually deal with hundred-thousand-level projections. For better rendering and interaction performance, we made the following technical choices. (i) **TC.1 Using WebGL instead of SVG as the front-end rendering technology to render the geospatial fossil collections.** The SVG layout is more popular in the visualization and visual analytics communities due to its superior interaction and convenient customization capabilities. However, the performance degrades rapidly when rendering thousands-level data because it adds multiple HTML nodes to visualize the projections on the map, and those nodes need to be rearranged for each interaction. WebGL renders data visualizations in a canvas node and data as pixel images. WebGL-based interactions refresh the image frames, and all operations are done within a single canvas node. Thus WebGL is a better choice for visualizing large-scale data and has already shown its effectiveness in scientific applications such as computer-aided drug design [21], protein structure analysis [11] and material microstructure characterization [12]. In our implementation, we utilize WebGL (over SVG) for rendering millions of fossil collections. (ii) **TC.2 Using the SVG layout to implement the fossil lens/ geological timescale view for better interactions.** This decision involved utilizing the excellent interaction ability supported by the SVG layout. Fossil lens determines whether the detected data are in the detection range of the lens by traversing the GPS in the data and updating the circular tree map of the lens with the user’s interaction.

## 4 EVALUATION AND DISCUSSION

We designed experiments to evaluate the scaling performance of `fossil-explorer.com`.

### 4.1 Metrics and Results

We focused on the interaction performance of large-scale data interactions. We simulated the actual operation process and identified two operations as key for impacting interaction performance: *t_zooming_* and *t_filtering_*. Based on this, we designed an interaction flow involving triple zoom in, triple zoom out,and a branch of the tree map to click on from the root node to the leaf node for data and timescale filtering. Each interaction flow was measured *n* times and the average interaction time taken to represent the performance of the interaction. Interaction performance was calculated by the following formula:

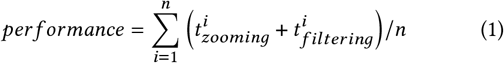

Since `fossil-explorer.com` is currently a static web application, we used the recorder provided by the developer tools in Chrome to record user interaction flows. The recorder is a preview feature in Chrome 97. It helps record, replay and measure user interaction flows. When we measured the performance of the user interaction flows, we set a listener event that waited for the page elements to finish loading. Interaction performance was measured only when all elements of the page were loaded. We recorded the performance of the interactions with different amounts of data and collected the script execution time during the interaction. As shown in Figure 3, we plotted the log-log performance curve of the system. The x-axis represents the amount of data and the y-axis the script execution time in milliseconds. It is clear that `fossil-explorer.com` achieved *sublinear scaling performance*, which is promising.

We also tested the interaction performance of the PBDB and GBDB. However, these two systems have backend data queries and graphic rendering available when interacting, and the recorder provided by Chrome did not measure query response time. Waiting too long for the graphics to render can also cause the recorder to time out and thus exit the measurement process. Therefore, we *manually measured* the interaction time of the systems and plotted their values in Figure 3, with the same log-log plotting. The figure shows that even when `fossil-explorer.com` loaded four times the amount of data as PBDB, the system exhibited better interactive performance. It is also worth noting that PBDB and GBDB aggregate and visualize data as circles to improve interactivity (using the SVG layout) and reduce the rendering burden on the browser. In contrast the rendering of large-scale data visualization technology utilizes SVG. This adds tens of thousands of HTML nodes to the page, resulting in longer interaction delays. Our results illustrate optimal performance for millions-level fossil data visualization. We believe that the combination of WebGL-based visualization and SVG-based interaction is a reasonable attempt to solve the challenges of large-scale data visualization.

### 4.2 Backend and Performance Impact

The current version of `fossil-explorer.com` is a static web application (i.e., the data are stored as static files in the CSV format). Here, we noted that the loading performance of the web application was similar for both static data file storage and using a database management system for the backend. However, the latter offers better run time performance, due to the following factors. (i) **Performance improvement from avoiding local file parsing.** A backend server (programmed using PHP or JavaScript) could efficiently convert query data from structured to JSON and transfer it to the front end using blocks. A file format such as CSV would also convert to JSON locally (though not as efficiently) before it could be utilized by the front-end application. (ii) **Performance improvement in the filtering operation.** Filtering operations such as timescale selection are some of the most frequently used functions in web applications. The front end could more quickly retrieve data from the database than directly filtering from the files as a database system support index. The improvement in performance will be more significant in larger datasets.

#### Suggested best practice

We suggest that anyone who wishes to deploy our tool should choose a proper backend server, including Nginx, Apache, Python, and others. However, we recommend using node.js to create the backend server and MongoDB to store the data. These can be well-adapted to the JavaScript-based front-end interface. MongoDB’s geospatial index can help with efficiently performing spatial queries on unions containing geospatial shapes and point sets, providing high-performance data querying and filtering functions.

## 5 RELATED WORKS

Databases and visualization systems gained increased popularity for domain paleotologic research over these years [18]. Most website front ends are built using JavaScript and a package of D3.js, and Leaflet.js. Many are developed to provide spatiotemporal information as the starting point. Examples include PBDB [17] and GBDB [19]. Other fossil database systems use a multimedia gallery as the entrance.

The advanced visualization challenges and analytics questions involved have also attracted interest from the computing community. Early works include using geospatial temporal visualization techniques for exploring biologic record distributions [2]. Novel visualization layouts such as TaxonTre and DoubleTree [9, 10, 16] have been proposed to address taxonomic name/location scaling performance and effective interactive qualities. Coordinated multiviews with interactive analytics loops have also been introduced to help answer complex scientific questions such as those regarding data quality [6], and assess whether they are suitable for studying biodiversity [16] and the relative habitat preferences between species over time within a particular region [3].

## 6 CONCLUSION

This research reports the ongoing work on `fossisl-explorer.com`, a sublinear interaction performance fossil data visualization approach. It currently features the multimedia graptolite fossil data, and the enhanced version also facilitates http://deepbone.org, the world’s most comprehensive database of vertebrate paleontology. The code is to be open access and we will continue to improve it.

## Footnotes

1 For example, https://fossiilid.info

